# Enhanced Complement Expression in the Tumor Microenvironment Following Neoadjuvant Therapy: Implications for Immunomodulation and Survival in Pancreatic Ductal Adenocarcinoma

**DOI:** 10.1101/2023.10.26.564099

**Authors:** Xiaofei Zhang, Ruoxin Lan, Yongjun Liu, Venu G Pillarisetty, Danting Li, Chaohui L Zhao, Suparna A. Sarkar, Weiguo Liu, Iman Hanna, Mala Gupta, Cristina Hajdu, Jonathan Melamed, Jessica Widmer, John Allendorf, Yao-Zhong Liu

**Affiliations:** Department of Pathology and Laboratory Medicine, New York University Grossman Long Island School of Medicine, Long Island, NY; Department of Biostatistics and Data Science, Tulane University School of Public Health and Tropical Medicine, New Orleans, LA; Department of Laboratory Medicine and Pathology, University of Washington School of Medicine, Seattle, WA; Department of Surgery, University of Washington School of Medicine, Seattle, WA; Department of Pathology and Laboratory Medicine, New York University Grossman School of Medicine, New York, NY; Department of Gastroenterology, New York University Grossman Long Island School of Medicine, Long Island, NY; Department of Surgery, New York University Grossman Long Island School of Medicine, Long Island, NY

**Keywords:** Pancreatic ductal adenocarcinoma (PDAC), Neoadjuvant therapy (NAT), Tumor microenvironment (TME), Complement, Immune exhaustion, Spatial Transcriptomics

## Abstract

**Purpose:** Neoadjuvant therapy (NAT) is increasingly being used for pancreatic ductal adenocarcinoma (PDAC) treatment. However, its distinct effects on carcinoma cells and the tumor microenvironment (TME) are not fully understood. This study employs spatial transcriptomics and single-cell RNA sequencing to investigate how NAT differentially remodels PDAC’s carcinoma cells and TME.

**Experimental Design:** We used spatial transcriptomics to compare gene expression profiles in carcinoma cells and the TME between NAT-treated and NAT-naïve PDAC patients and correlated with their clinicopathologic features. Complementary single-nucleus RNA sequencing (snRNA-seq) analysis was conducted to validate our findings and identify cell types driving NAT-induced gene expression alterations.

**Results:** We found NAT not only induces apoptosis and inhibits proliferation in carcinoma cells but also significantly remodels the TME. Notably, NAT induces a coordinated upregulation of multiple key complement genes (C3, C1S, C1R, C4B and C7) in the TME, making the complement pathway one of the most significantly affected pathways by NAT. Patients with higher TME complement expression following NAT exhibit improved overall survival; more immunomodulatory and neurotrophic cancer-associated fibroblasts (CAFs); more CD4^+^ T cells, monocytes and mast cells; and lower immune exhaustion gene expression. snRNA-seq analysis demonstrates C3 complement upregulation specifically in CAFs but not in other stroma cell types.

**Conclusions:** Our findings indicate that NAT may reduce immunosuppression in PDAC by enhancing complement production and signaling within the TME. These findings suggest that local complement dynamics could serve as a novel biomarker for prognosis, evaluating treatment response and resistance, and guiding therapeutic strategies in NAT-treated PDAC patients.

**Translational Relevance:** As neoadjuvant therapy (NAT) increasingly becomes the preferred approach in treating resectable and borderline resectable pancreatic ductal adenocarcinoma (PDAC), there is a growing demand for novel biomarkers specifically tailored for post-NAT PDAC patients. Our study focused on how NAT differentially remodels the tumor cells and the tumor microenvironment (TME). We demonstrate that NAT can enhance local complement production and signaling in PDAC’s TME, which is associated with reduced immune exhaustion and improved overall survival. Our results highlight the importance of local complement dynamics in influencing treatment response, resistance, and overall clinical outcomes in PDAC. This new mechanism provides new opportunities in biomarker development which could facilitate more accurate prognostication and precise treatment stratification, particularly for patients undergoing NAT in PDAC.

## INTRODUCTION

Pancreatic ductal adenocarcinoma (PDAC) is a highly aggressive cancer with a five-year survival rate of only 11% (1). It is projected to become the second leading cause of cancer-related deaths in the United States by 2030(2). Only 15-20% of patients are eligible for surgery, as most patients present with locally advanced unresectable disease or distant metastases(3). Furthermore, since most patients who undergo surgical resection ultimately succumb to their disease, systemic chemotherapy is a necessary therapeutic component for most patients(4).

PDAC presents a significant therapeutic challenge due to its resistance to various treatment modalities including immune checkpoint inhibitor therapy(5,6). It exhibits extensive genetic diversity, leading to the development of subpopulations with diverse molecular alterations and resistance mechanisms(7). The tumor microenvironment (TME) in PDAC is highly immunosuppressive, characterized by the presence of immunosuppressive cells, cytokines, and chemokines that impede effective anti-tumor immune responses despite the presence of large numbers of effector T cells(8–10). Additionally, the TME plays a pivotal role in influencing tumor growth and metastasis through its impact on mitochondrial biogenesis, blood supply, and nutrient availability (11). These factors collectively contribute to the high resistance observed in PDAC, highlighting the need for innovative combinational strategies to improve treatment outcomes.

Neoadjuvant therapy (NAT) has emerged as the preferred approach for managing resectable and borderline resectable PDAC over adjuvant therapy for several reasons. Firstly, NAT offers better tolerability for patients by administering chemotherapy and/or radiation prior to surgery, taking advantage of patients’ relatively better health status and minimizing treatment-related complications(12,13). Secondly, NAT effectively increases tumor resectability and R0 resection rate by downstaging locally advanced disease and facilitating complete tumor removal(6,14). Moreover, NAT targets early metastasis, potentially improving treatment outcomes(6,14). It also provides an opportunity to observe an individual patient’s response to therapy for selecting suitable candidates for surgery(6,14). Recently, some studies suggested that NAT may be associated with improved overall survival rates compared to adjuvant therapy (12,13). Furthermore, recent studies suggested that NAT may have the potential to prime the stroma and enhance the response to immunotherapy in various cancers (15,16). By modulating TME, NAT may render PDAC more susceptible to immunotherapeutic interventions (15,16).

Single-cell RNA sequencing (scRNA-seq) is a valuable tool for characterizing TME as it provides insights into cellular heterogeneity and molecular subtypes(17,18). Several scRNA-seq studies have explored the PDAC TME, offering valuable insights into this complex ecosystem(19–35). However, scRNA-seq techniques lack spatial context, and are susceptible to cell dissociation-associated artifacts. Spatial transcriptomic analysis complements and extends the scope of single-cell studies (36,37) by providing valuable insights into the spatial distribution of gene expression and a holistic characterization of the PDAC TME. To obtain a comprehensive view of the PDAC TME and its response to NAT, we performed a spatial transcriptomic comparative study on formalin-fixed paraffin-embedded (FFPE) specimens from PDAC patients who received NAT before surgery versus those who underwent upfront surgery without NAT.

## Materials and Methods

### Patient cohort and data collection

All patient selection and data collection procedures were reviewed and approved by the University of Wisconsin – Madison Institutional Review Board. The inclusion criteria were patients with non-metastatic PDACs who had surgery at the University of Wisconsin–Madison Hospital between 2015 to 2020. Patients with incomplete clinicopathologic information or incomplete archived slides were excluded. A total of 36 patients met these criteria and were used for this study, including 13 patients who had upfront surgery without any forms of NAT and 23 patients who had NAT (either chemotherapy alone or in combination with radiation) before surgery. All patients had at least 38 months of clinical follow-up. The pertinent clinicopathological information, including age, gender, carcinoma size, regime of NAT, type of surgery, pathological grade and stage, and treatment response, were retrospectively collected by reviewing electronic medical records. Among the 23 patients in the NAT group, 16 patients received multiple cycles FOLFORINOX, and 7 patients received gemcitabine and Abraxane. Eight patients received concurrent radiation therapy. For all patients, only resection specimens were used, except for 4 patients, for whom the pre-NAT biopsies were also included for a paired comparison with their corresponding post-NAT resection samples.

### Sample preparation for NanoString GeoMx DSP

Serial sections (4 μm in thickness) were cut from the selected formalin-fixed, paraffin-embedded (FFPE) tissue block for the consecutive H&E staining, immunofluorescence staining for DSP (Digital Spatial Profiler) experiment. For DSP experiment, sections were baked at 60⁰C for 1.5 hours and deparaffinized in CitriSolv. After rehydration, sections were incubated in 100 °C 1xTris-EDTA/pH 9 buffer for 15 minutes for antigen retrieval. After digestion and post-fixation, all sections were hybridized to UV-photocleavable barcode-conjugated RNA in situ hybridization probe set (Nanostring CTA, 1812 gene targets) overnight at 37 °C. Then, all slides were washed to remove off-target probes. Next, slides were used for immunofluorescence staining with morphology markers set, which contains 1:10 SYTO13 (Thermo Fisher Scientific, cat. no. 57575), 1:20 anti-panCK-Alexa Fluor 532 (clone AE-1/AE-3; Novus Biologicals, cat. no. NBP2-33200AF532), and 1:100 anti-αSMA-Alexa Fluor 647 (clone 1A4; Novus Biologicals, cat. no. IC1420R) in blocking buffer (NanoString). For each slide, 2–5 ROIs (Regions of Interest) with the most representative morphology were selected. Each ROI was segmented into a carcinoma AOI (area of illumination, panCK+, SMA-) and aTME AOI (panCK-, SMA+) using the scanned the immunofluorescence image. Photocleavage of spatially indexed barcode and sample collection were done on GeoMx DSP equipment (NanoString). Samples obtained from each AOI were collected in a 96-well plate. RNA Libraries were prepared according to the manufacturer’s instruction.

Then, Next Generation Sequencing (NGS) read-out was done using an Illumina NextSeq 550AR sequencer. The data was exported as digital count conversion (DCC) files which were used for analysis using R and Bioconductor packages.

### Data pre-processing and normalization

We followed a published online R and Bioconductor GeoMx-NGS workflow (NanoString Technologies, Inc) for QC (quality control), pre-processing and normalization of raw GeoMx DSP count files and metadata. The website of the workflow is at https://bioconductor.org/packages/devel/workflows/vignettes/GeoMxWorkflows/inst/doc/GeomxTools_RNA-NGS_Analysis.html. In brief, the workflow included data loading (of DCC, PKC and annotation files) and illustration of study design using Sankey diagram (Supplemental Figure 1). The loaded data then underwent QC and pre-processing steps, including segment QC for every AOI segment (e.g., removing segments with >1,000 raw reads and those with less than 80% rate for % aligned, % trimmed or % stitched), probe QC to remove low-performing probes, creation of gene-level count data by combining counts across multiple probes of a gene via geometric mean, and determination of limit of quantification (LOQ) of each segment based on the distribution of negative control probes. Based on the calculated LOQ, the number of detected genes was decided for each segment and segment-wise gene detection rate was inferred, and those segments with less than 1% of gene detection rate were removed. Similarly, the detection rate for each gene across the study was determined and those genes detected in only 5% or less of all segments were removed. Based on the filtered gene expression matrix, the GeoMx data were normalized using Q3 (3^rd^ quartile) normalization. The Q3 normalized data were then used for downstream data analyses.

### Unsupervised analysis

The UMAP method was used to visualize the segment-level data in two major dimensions (UMAP1 and UMAP2) by using the UMAP R package. Unsupervised hierarchical clustering of all genes was performed using pheatmap package by first conducting log2 transformation of the data, followed by selecting highly variable genes (i.e., those of top 20 percentile of coefficient of variation of gene expression) to be included in the cluster heatmap. Similarly, unsupervised hierarchical clustering of 5 complement genes (C3, C1S, C1R, C4B and C7) was performed with the pheatmap package using subject-wise average of gene expression value in the TME.

### Differential expression analysis

Linear modeling analysis implemented in R was conducted to detect the effect of NAT on gene expression profiles in either the carcinoma cells AOIs or the TME AOIs. Specifically, the log2 transformed subject-wise average of gene expression of carcinoma AOIs or the TME AOIs was modeled as the dependent variable, with independent variables including treatment status (i.e., NAT vs. naïve), age, gender, pathology grade and stage of the tumor. The estimate and the p value for the regression coefficient for treatment status (NAT-treated vs. naïve) were used to represent the effect of differential expression induced by NAT. To illustrate the differential expression, volcano plots were made with EnhancedVolcano package (version 1.13.2).

### Signature scores for immune and fibroblast cell types

Signature scores for immune cell types of each AOI were imputed from the gene expression matrix with SpatialDecon package (version 1.4.3). Scores for other signature scores, including Malignant State Programs (i.e., cycling (S), cycling (G2/M), MYC signaling, adhesive, ribosomal, interferon signaling and TNF-NFkB signaling), Malignant Lineage Program (i.e., acinar-like, classical-like, basaloid, squamoid, mesenchymal, neuroendocrine-like, neural-like progenitor) and Fibroblast Programs (i.e., adhesive, immunomodulatory, myofibroblastic and neurotropic), were inferred based on geometric average of signature genes as described by Hwang et al. in 2022 (24). The signature score for Immune Exhaustion was inferred from the signature genes as provided in Werba G et al., in 2023 (33). The same linear modeling analysis was used to test the effect of NAT on the above inferred signature scores.

### Gene set enrichment analysis

The differentially expressed genes with p value < 0.05 were submitted to DAVID enrichment analysis (38,39) portal (https://david.ncifcrf.gov/home.jsp) for gene set enrichment analysis. To infer up-or downregulated gene sets, the up-or downregulated genes were submitted for analysis separately. To visualize the enriched gene sets, dot plots were made with ggplot2 R package (v3.4.2) and chord maps were plotted with GOplot package (v1.0.2).

### Survival analysis

R packages, survival (v3.5.5) and survminer (v0.4.9), were used to make Kaplan Meier (KM) survival curve and compare and test significance for the difference of survival between groups of patients. Specifically, the Cox Proportional Hazards Model was used to test the significance of survival differences in groups of patients while controlling for age, gender, tumor size and cancer grading.

### Single cell nucleus RNA-seq data analyses

Single-nucleus RNA-seq (snRNA-seq) data of Hwang et al. (24) was downloaded from Single Cell Portal at https://singlecell.broadinstitute.org/single_cell/study/SCP1089 (for 88031 naïve cells from 15 NAT-naïve PDAC patients) and https://singlecell.broadinstitute.org/single_cell/study/SCP1096 (for 50516 treated cells in 11 NAT-treated patients). Specifically, for the purpose of the analyses in the work herein, we only downloaded the snRNA-seq data of the already annotated fibroblasts, carcinoma cells, and immune cells for untreated (naïve) and treated (NAT) patients.

We used Seurat pipeline (40,41) for the initial processing and analyses of the downloaded data. The major analyses involved data merging of treated and naïve dataset, data normalization, identification of 2,000 most variable genes, dimensionality reduction and visualization of gene expression in UMAP plots, and within UMAP plots, highlighting and comparing between treated and naïve groups for expression of important genes, such as C3, in specific cell populations.

To annotate and define subtypes of CAFs, signature genes of each specific groups as described in Hwang et al. (24) were used. Assignment of CAFs to a specific subtype follows the method in Hwang et al. (24). Specifically, if a cell’s signature gene score for a subtype is higher than the cell’s background gene score, the cell will be assigned to the subtype. For a given cell, the signature or background gene score was calculated via the mean of log2 transformation of the associated list of genes’ expression (GE) values. The background gene set was determined by selecting those genes that matched with individual signature genes in terms of ranks of the across-cell mean expression (i.e., the mean log2 expression) determined for all expressed genes in CAFs.

A linear mixed model was used to compare a gene’s expression in NAT-treated vs. naïve patients via lme4 package in R. The model was constructed as: [log2(GE) ∼ treatment + (1|subject)], which accounted for subject-level random effects. Similarly, using MASS library in R, a generalized linear mixed model was used to compare the proportion of C3 high-expression cells in treated vs. naïve patients. The model was constructed as: [C3 high-expression cells ∼ Treatment, random = ∼1|subject, family = binomial]. Here, a C3 high-expression cell was defined as a cell, whose C3 expression is above the median of non-zero counts of all cells. A similar approach with generalized linear mixed model was taken to compare treated vs. naïve patients for the proportion of immune cells with a high expression of immune exhaustion genes (CD274, CTLA4, HAVCR2, LAG3 and TIGIT), with median of non-zero counts as the cutoff to define the cells of high expression for the above genes. For the above analyses, one tailed p values were used since the analyses were based on the hypotheses that there is increased complement gene expression in fibroblasts and increased fibroblasts with a high expression of complement genes in NAT vs. naïve patients. Within immune cells, our hypothesis is that there is a decreased expression of immune exhaustion genes and decreased immune cells with a lower expression of immune exhaustion genes in NAT-treated vs. naïve patients.

### Data availability

The spatial transcriptomics data used here along with the metadata were submitted to NCBI GEO (Gene Expression Omnibus) with an accession number of GSE240078.

## RESULTS

### Intra- and intertumoral heterogeneity of NAT response in PDAC carcinoma and TME

The clinicopathologic characteristics of the 36 patients in our study cohort are summarized in Figure 1A and Supplementary Table 1. This cohort includes 13 patients who had upfront surgery (the naïve group) and 23 age-, gender-, and stage-matched patients who had surgery after receiving neoadjuvant chemotherapy (the NAT group). The NAT consisted of either FOLFIRINOX or gemcitabine treatment, with or without radiation. Only resected primary PDAC specimens were used in this study except for 4 patients whose pre-NAT biopsies were included in the naïve group and post-NAT resection specimens were included in the NAT group. A separate pairwise longitudinal comparison of the pre-NAT biopsy versus the post-NAT resection from the same patient was also done for these 4 patients (Figure 1E and Supplementary Tables 2). To assess the relative extents of intra-vs. intertumoral heterogeneity, 2 to 5 ROIs (region of interest) were selected from each case, amounting to a total of 119 ROIs from the 36 patients in the entire cohort (Supplementary Figure 1). Using immunofluorescence staining, each ROI was segmented into two AOIs (Area of Illumination): the carcinoma AOI (pan-CK+, SMA-) containing only malignant cells, and the TME AOI (pan-CK-, SMA+) which comprised of various types of stroma cells and immune cells (Figure 1B). RNA libraries were prepared from these 238 AOIs, of which 223 (93.7%) passed the quality control and were used for analysis, including 42 carcinoma AOIs and 43 TME AOIs from the naïve group, and 72 carcinoma AOIs and 66 TME AOIs from the NAT group.

**Figure. 1.**
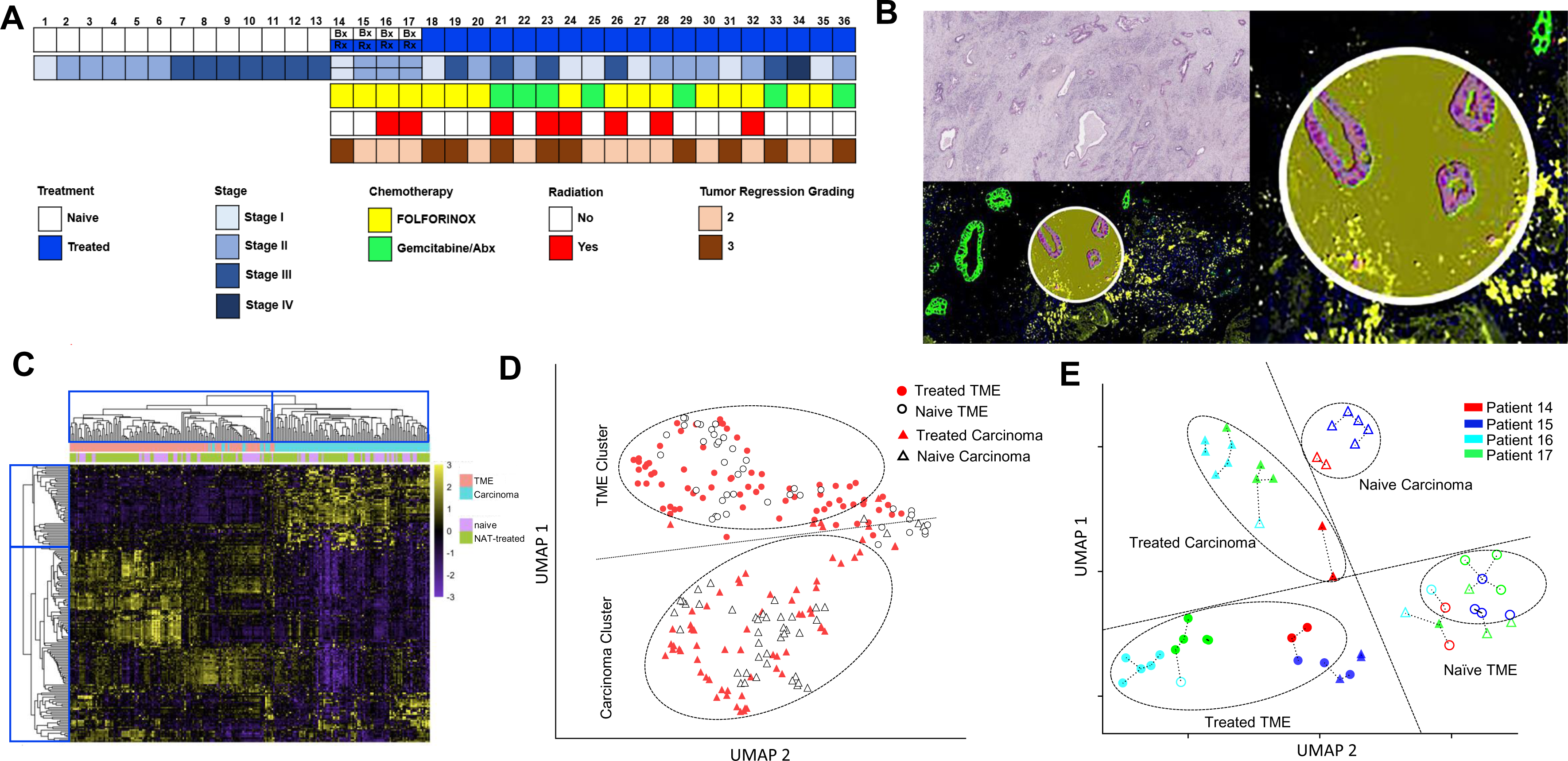
Intra- and intertumoral heterogeneity of NAT response in PDAC carcinoma cells and TME revealed by spatial transcriptomics. **A.** Clinical and pathologic characteristics of the 36 patients in our study cohort. For patients 14, 15, 16 and 17, pre-NAT biopsies were included in the naïve group, and the post-NAT resection specimens were included in the NAT group. For all other patients, only resected primary PDAC specimens were used. **B.** Immunofluorescence segmentation of each ROI into carcinoma AOI and TME AOI (pan-CK in green, α-SMA in yellow, DAPI in blue). Total mRNA libraries were prepared separately from each of these two AOIs for every ROI. **C.** Heatmap of gene expression profiles in all 223 carcinoma and TME AOIs. Distinct separation between carcinoma AOIs and TME AOIs is evident in the unsupervised clustering analysis, with only a limited overlap. There is no discernible separation between the naïve group and the treated group in either the carcinoma AOI cluster or the TME AOI cluster. **D.** UMAP plot demonstrates the near-complete separation of carcinoma AOIs (below the dashed line) and TME AOIs (above the dashed line). 95.1% (137/144) of the TME AOIs cluster above the dashed line, and 94.5% (103/109) of the carcinoma AOIs cluster below the dashed line. Approximately 5% of the AOIs overlap at the peripheries of these two clusters. There is no noticeable separation between the NAT group and the naïve group within either cluster. **E.** UMAP plot of the four pairs of pre-NAT biopsy (as naïve) and post-NAT resection (as NAT-treated) in patients 14, 15, 16, 17 (Δ=naïve carcinoma; ▴=NAT-treated carcinoma. ○=naïve TME; ●=NAT-treated TME. Datapoints from same patient are shown in the same color). For both carcinoma AOIs and TME AOIs, datapoints from NAT group are separated from those from the naïve group. The dashed lines connect each datapoint to its nearest neighbor. Most of the datapoints are closer to datapoint from same patient than to those of the different patients, suggesting datapoints from same patient tend to aggregate together.

First, we compared the gene expression profiles between carcinoma AOIs versus TME AOIs. Carcinoma AOIs showed significantly higher expression of 302 and 321 genes than TME AOIs in the naïve group and the NAT group, respectively, with an overlap of 261 genes which included *KRT19, KRT7, KRT18, MUC1, CDH1, EPCAM, JUP, ITGA3, ITGB4, ERRB2, LAMB3, CAPN2, LGALS3, LCN2 etc*. In contrast, TME AOIs showed significantly higher expression of 286 and 294 genes than carcinoma AOIs in the naïve group and the NAT group, respectively, with an overlap of 250 genes including *PDGFRB, COL6A3, FCGR2A/B, IL1R1, etc.* (Supplementary Figure 2, Supplement Data 1). Hierarchical clustering analysis successfully grouped these 223 AOIs into the carcinoma cluster and the TME cluster, with a limited number of datapoints appearing in between these two distinct clusters. There is no discernible differentiation between the naïve and NAT AOIs within both the carcinoma AOI cluster and TME AOI cluster (Figure 1C).

Next, we used a dimension reduction technique to visualize the clustering pattern of these 223 AOIs in two major UMAP dimensions based on their transcriptomics profile. Like the results from the hierarchical clustering analysis, these 223 AOIs were largely separated into carcinoma cluster and the TME cluster, with only a limited number of AOIs from each category intermingling at the peripheries of these two clusters (Figure 1D). However, notable differentiation based on treatment status (i.e., naïve vs. NAT) was once again not evident, suggesting both carcinoma cells and the TME responded to NAT heterogeneously at the transcriptomics level.

To further investigate the roles of intra-versus intertumoral heterogeneity in NAT response, we also made a UMAP plot of the four pairs of samples in which the pre-NAT biopsy was compared with the post-NAT resection specimen from same patient. In addition to the clear separation of carcinoma AOIs from TME AOIs, a near-complete separation was also noted between the naïve group (i.e., pre-NAT biopsy) and the NAT group (i.e., the post-NAT resection) in both carcinoma AOI cluster and TME AOI cluster (Figure 1E). In accordance with the anticipated relative magnitudes of intra-versus intertumoral heterogeneity, the AOIs from the same patient tended to aggregate more closely together than those from different patients (Figure 1E).

To compare the cellular composition within carcinoma AOIs and TME AOIs, we calculated the gene signature scores for different types of immune cells and fibroblasts using our transcriptomics data. As compared to carcinoma AOIs, TME AOIs contained significantly more fibroblasts and immune cells across the board (Supplementary Figure 3).

### Effects of NAT on PDAC carcinoma cells

We studied the effect of NAT on PDAC carcinoma cells by comparing the gene expression profiles in the carcinoma AOIs between the naïve group versus the NAT group. After controlling for age, gender and other clinicopathological factors, we identified 44 NAT-induced differentially expressed genes (DEGs) in carcinoma AOIs, including 22 upregulated and 22 downregulated (Figure 2A, Supplementary Data 2). The most significantly upregulated genes include *DDB2* (p = 7.70e-4), *FBP1* (p= 0.0016) and *PRKACB* (p = 0.0028). The most significantly downregulated genes include *IL32* (p =0.0039), *ACTB* (p = 0.0041) and *TPM1* (p = 0.0059). DAVID enrichment analysis of these DEGs indicated that the upregulated genes in the carcinoma cells of NAT-treated PDAC were significantly enriched with functional terms related with apoptosis (KW-0053∼Apoptosis, FDR = 0.013) and Ser/Thr-type protein kinases (SM00133:S_TK_X, FDR = 0.024), suggesting upregulation of these functional terms by NAT (Figure 2B). On the other hand, the downregulated genes were significantly enriched with functional terms related to chromatin (GO:0000785∼chromatin, FDR = 0.0059), nucleoplasm (GO:0005654∼nucleoplasm, FDR = 0.023), GO:0032991∼macromolecular complex (FDR = 0.023), and KW-0488∼Methylation (FDR = 0.024) (Figure 2B), suggesting downregulation of these functional terms by NAT. These results are consistent with the expected cytoreductive effect of neoadjuvant chemotherapy on carcinoma cells.

**Figure. 2.**
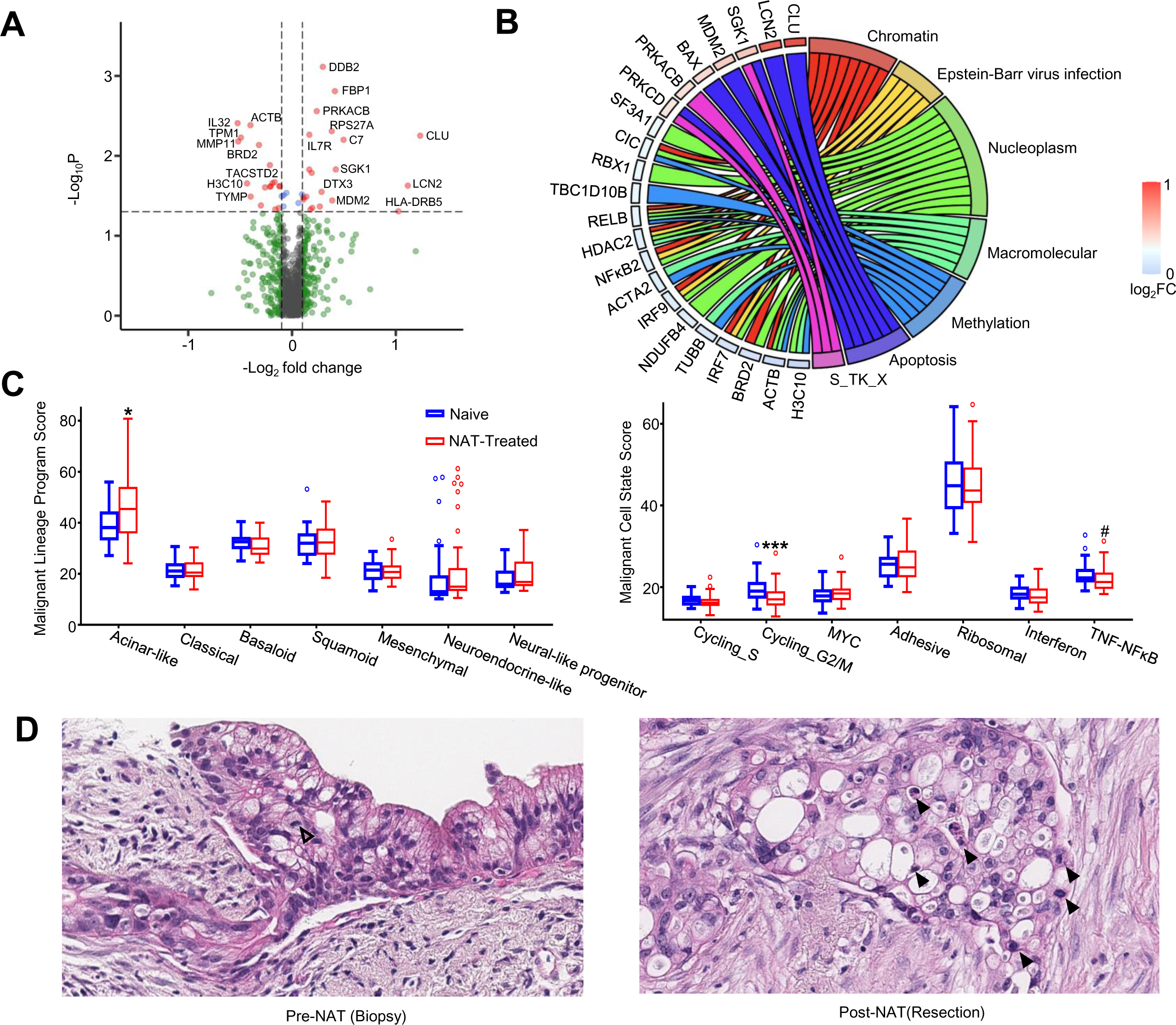
Impact of NAT on PDAC carcinoma cells. **A.** Volcano-plot showing up- and down-regulated genes in carcinoma cells induced by NAT. **B.** Chord map illustrating the relationship between individual DEGs and the enriched pathways. **C.** Impact of NAT on malignant cell lineage programs (left) and malignant cell states (right). NAT treatment significantly upregulated the acinar program and downregulated the cycling_G2/M state. TNF-NFκB signaling is also decreased with marginal statistical significance (* p<0.05; *** p<0.001, # p<0.1). **D.** Histologic evaluation confirming that the NAT-treated carcinoma cells show significantly increased apoptosis (solid arrowhead), decreased mitosis (open arrowhead), greater nuclear pleomorphism, and cytoplasmic vacuolization, in line with the transcriptomics findings.

Next, we investigated the impact of NAT on carcinoma cells in more detail by utilizing the recently defined gene signatures for the seven malignant cell programs representing the lineages of carcinoma cells and the seven malignant cell states^19^. Signature scores for each of these malignant programs and cell states were calculated for all carcinoma AOIs based on the geometric mean of expression levels for their respective signature genes set. Their association with NAT was assessed after controlling for age, gender, cancer grade and stage. We found that there was a significant increase of acinar-like program (p = 0.011, Figure 2C) and a significant decrease of cyclining_G2/M state (p = 0.001), along with a marginally significant decrease of TNF-NFkB signaling (p = 0.057) in the NAT group. These findings were also supported by our morphologic analysis, which showed increased apoptosis along with decreased proliferation and mitosis in the carcinoma cells of the NAT group compared to the naïve group (Figure 2D).

### Effects of NAT on the TME of PDAC

We examined the effect of NAT on PDAC TME by comparing the gene expression profiles of the TME AOIs between the NAT versus the naive groups. After controlling for age, gender, carcinoma grade and stage, we identified 52 NAT-induced DEGs in the TME AOIs, including 28 upregulated and 24 downregulated (Figure 3A, Supplementary Data 3). The top significantly upregulated genes include *CDKN1A* (p=3.82e-4)*, MDM2* (p=8.04e-4)*, FAS* (p = 2.72e-3) and *IL6ST* (p = 0.0055). The top significantly downregulated genes include *HDAC2* (p = 2.84e-5), *TPM1*(p=7.21e-4), *ACTB* (p = 2.28e-3) and *PKM* (p = 0.0033). Remarkably, there were five complement genes (*C1S, C1R, C3, C4B* and *C7*) among the significantly upregulated genes, suggesting that NAT induced a coordinated upregulation of complement pathway genes in the TME of PDAC.

**Figure. 3.**
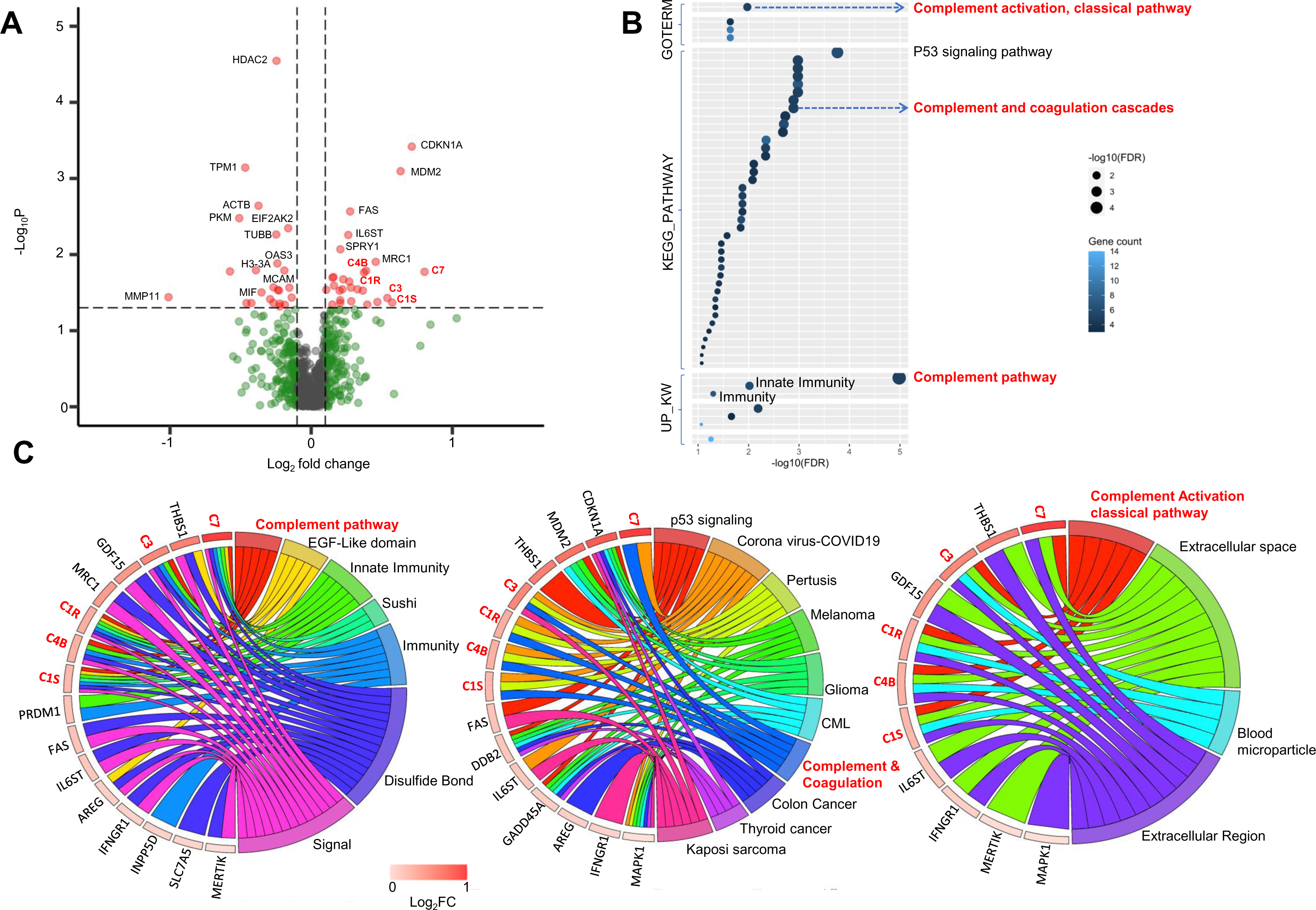
Impact of NAT on PDAC TME. **A.** Volcano plot showing the significantly up- and downregulated DEGs in TME induced by NAT. **B.** Dot Plots summarizing the most upregulated pathways in TME following NAT. Three complement related pathways (highlighted in red) were significantly upregulated. **C.** Chord maps illustrating the relationship between individual DEGs and the enriched pathways. The five NAT-upregulated complement genes are broadly associated with regulation of immune related pathways.

DAVID enrichment analysis of these significantly upregulated DEGs showed that NAT upregulated more functional terms in the TME than in the carcinoma cells of PDAC (Figure 3B). Among those, there were three highly enriched terms related to complement pathway, including KW-0180∼Complement pathway (FDR = 1.04e-5), GO:0006958∼Complement activation, classical pathway (FDR = 1.06e-2), and has04610∼Complement and coagulation cascades (FDR = 1.28e-3). Additionally, there were functional terms related to immunity, including KW-0399 Innate immunity (FDR = 9.61e-3); KW-0391 Immunity (FDR = 5.04e-2), and multiple pathways related to cancer signaling such as hsa04115 p53 signaling pathway (FDR = 1.73e-4) (Figure. 3B). To illustrate the gene components for these enrichment terms, chord maps were made for the most significantly enriched GO, KEGG and UP_KW functional terms and their related genes. The 5 complement genes (C1S, C1R, C3, C4B and C7) contribute to not only complement-related pathways, but also other key functional terms related to innate and adaptive immunity (Figure 3C). These findings imply that NAT-induced upregulation of complement expression within TME may serve as a local regulatory hub in TME and contribute to the overall immunomodulation in PDAC.

To investigate how NAT influences the tumor immune microenvironment (TIME) in PDAC, signature scores for all major immune cell types were calculated for each TME AOI and tested for association with NAT treatment while controlling for age, gender, carcinoma grade and stage. There was no statistically significant change of signature scores for most types of immune cells in the TIME of the NAT group compared to the naïve group, except for mast cells which were increased with a marginal statistical significance (p=0.09) (Figure 4A). Next, we studied the effect of NAT on the cancer-associated fibroblasts (CAFs) within PDAC TME using the expanded CAF classification system recently defined by Hwang et al. in 2022^19^, in which CAFs are categorized into 4 specific programs based their gene expression profiles, including myofibroblastic CAFs, adhesive CAFs, immunomodulatory CAFs, and neurotrophic CAFs. We computed signature scores for all four programs of CAFs within each TME AOI and analyzed the association with NAT treatment while controlling for age, gender, carcinoma grade and stage. We found that immunomodulatory (p=0.023) and neurotropic CAFs (p=0.002) were significantly increased in the TME of the NAT group compared to the naïve group (Figure 4B).

**Figure 4.**
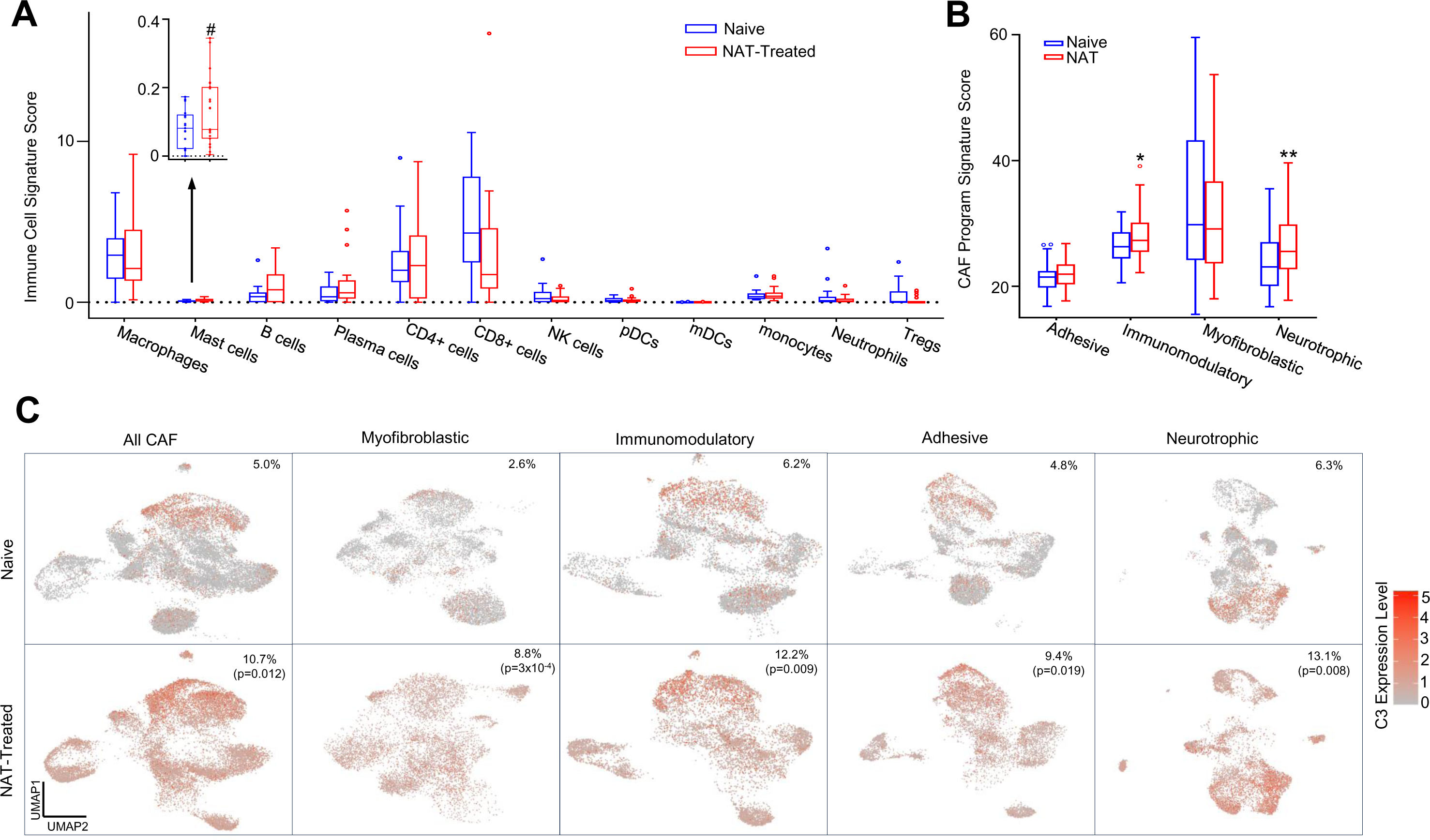
Impact of NAT on CAFs and immune cells in PDAC TME. **A.** Analysis of changes in immune cell composition in the tumor immune microenvironment (TIME) post-NAT. No statistically significant changes were observed except a marginal increase in the number of mast cells. **B.** NAT significantly increases the immunomodulatory CAF and the neurotrophic CAF, with no significant changes in other CAFs. **C.** snRNA-Seq analysis of data from Hwang et al. demonstrates that the number of complement C3 expressing CAFs was significantly increased following NAT. The percentages of C3 expressing CAFs and p-values are labeled on the upper right corner.

To identify which specific cell types in the TME are primarily responsible for the upregulation of complement genes following NAT, we analyzed a single-nucleus RNA (snRNA-seq) sequencing dataset from Hwang et al. (24), which was archived in the single cell portal. This dataset encompassed the gene expression profiles of 22,164 genes across 88031 cells from 15 NAT-naïve patients and 50516 cells from 11 NAT-treated patients. Our analysis revealed a significant increase of C3 expression in CAFs of the NAT-treated patients compared to NAT-naïve patients (p = 0.044). Based on generalized linear mixed model analysis, the proportion of C3 high expression CAFs was found to be significantly higher in NAT-treated patients than NAT-naïve patients, (10.66% vs. 5.03%, p = 0.012). (Figure 4C). In contrast, no notable changes in C3 expression level were observed in other types of cells including tumor cells and immune cells. A more detailed analysis on subtypes of CAFs showed that all subtypes of CAFs exhibited increased complement C3 gene expression, with the most pronounced increases were observed in immunomodulatory and myofibroblastic CAFs. These results indicate that the CAFs are the primary contributing factor of the NAT-induced upregulation of complement gene expression in TME.

### NAT-induced upregulation of TME complement expression is associated with improved survival and immunomodulation

To assess the clinical and prognostic significance of NAT-induced upregulation of TME complement, we divided our cohort’s NAT group into high and low complement subgroups. This stratification was based on clustering analysis of the mean expression levels of five complement genes upregulated by NAT (C3, C4, C7, C1S and C1R) in the TME. Out of 23 NAT-treated patients in our cohort, 9 were categorized into the high complement subgroup and 14 into the low complement subgroup (Figure 5A). Kaplan-Meier analysis indicated that the high complement subgroup demonstrated a significantly better overall survival compared to the low complement subgroup (p=0.023) (Figure. 5B). Additionally, compared to the naïve group, the high complement subgroup showed a marginally better survival trend (p<0.1), while the low complement subgroup’s survival rates were not significantly different (Figure. 5B). Multivariate Cox proportional hazard ratio analysis further identified a high TME complement level as an independent favorable prognostic factor (hazard ratio = 0.21; 95% Confidence Interval (CI): 0.06 to 0.81) for NAT-treated patient (Figure. 5C). In contrast, a positive surgical resection margin (R1) was recognized as an independent adverse prognostic factor (HR=5.45, 95% CI=1.28-23.27) (Figure. 5C).

**Figure. 5.**
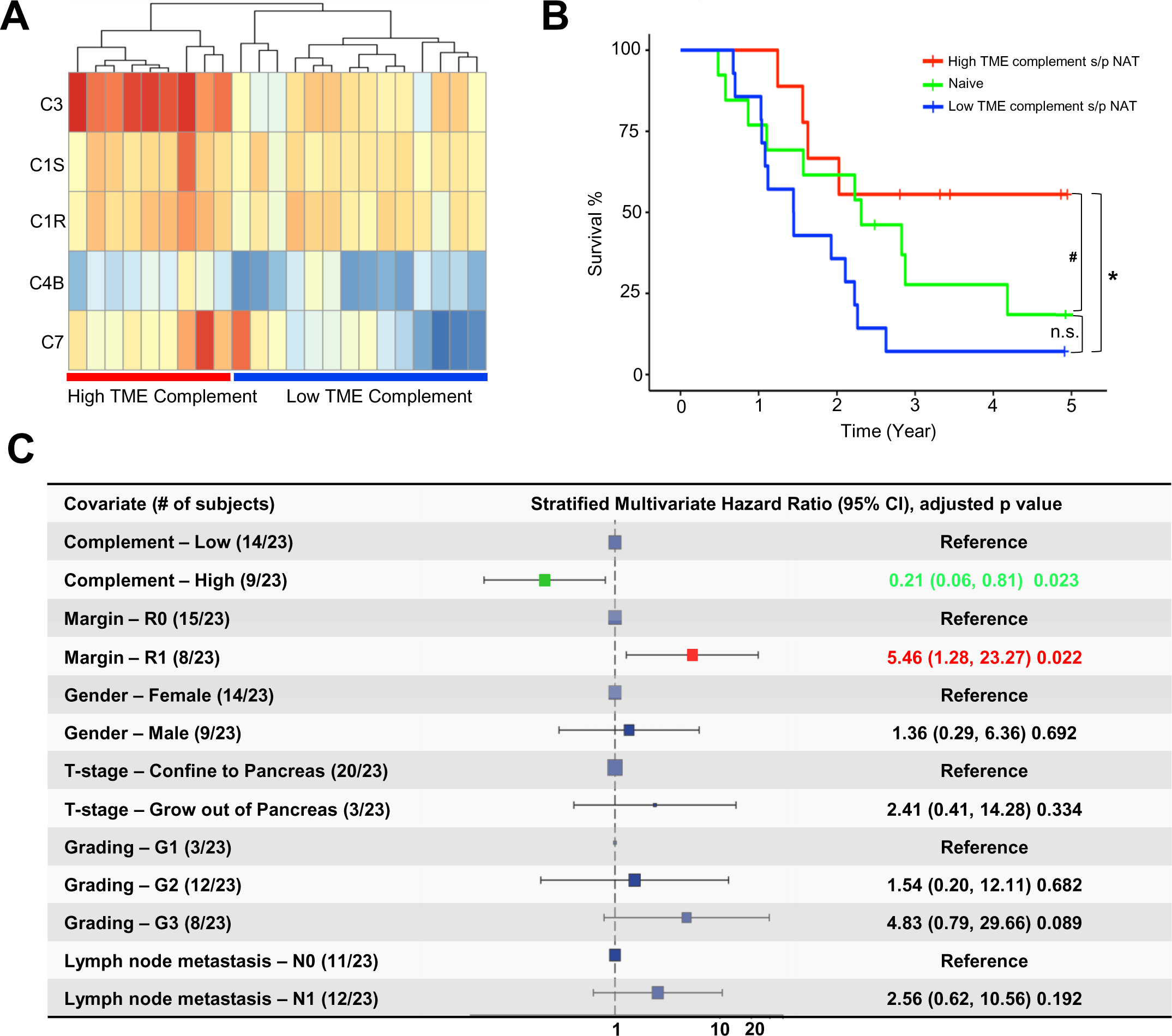
Prognostic relevance of NAT-induced upregulation of TME complement. **A.** Heatmap displaying the expression levels of the five key complement genes in the TME of the NAT group. Unsupervised clustering effectively categorized them into the high and low TME complement subgroups. **B.** Kaplan-Meier curve showing that the high TME complement subgroup (red line) has a significantly improved overall survival compared to the low TME complement subgroup (blue line, p=0.02) and a marginally improved overall survival compared to the naïve group (green line, p=0.09) (* p<0.05; # p<0.1). There is no statistically significant difference in overall survival between the low TME complement subgroup and the naïve group. **C.** Forest plot of multivariate Cox proportional hazards model identifying the TME complement level as an independent favorable prognostic factor for NAT-treated patients (HR=0.21; 95% CI: 0.06-0.51, p=0.023). Positive surgical resection margin (R1) is identified as an independent poor prognostic factor for NAT-treated patients (HR=5.46; 95% CI: 1.28-23.27, p=0.022).

To further clarify the link between the elevated complement level in TME and survival benefits in post-NAT patients, we also conducted a longitudinal paired analysis of complement C3 level in the TME of the four patients for whom we had both pre-NAT biopsy and post-NAT resection samples. We examined the changes in TME C3 levels and their correlation with clinical outcome. Among these 4 patients, two of them displayed a significantly increased TME C3 level following NAT and showed improved overall survival compared to the rest two patients who did not show increase of TME C3 level post-NAT (Supplementary Figure 6). These results imply that patients with a significant upregulation of TME complement level in response to NAT could be indicative of improved overall survival.

To elucidate the mechanism behind the association between the NAT-induced upregulation of TME complement and the observed improvement in patient survival, we compared the signature scores for all immune cells and CAF subtypes between the high and low complement subgroups. Our results showed that the high complement subgroup exhibited significantly higher signature scores for mast cell (p = 0.037) and CD4^+^ T cells (p = 0.039) compared to the low complement subgroup (Figure. 6A). Moreover, the high complement subgroup showed significantly higher signature scores for immunomodulatory and neurotrophic subtypes of CAFs (p = 3.46e-4, Figure 6B). In addition, the high TME complement subgroup also showed a notable reduction of expression for immune exhaustion gene signature (p = 0.038) while maintaining stable expression of immune cell cytotoxicity gene signature (Figure. 6B). Further analysis using snRNA-seq data from Hwang et al. (24) revealed reduced expression of immune checkpoint gene *CD274* (p=0.019) and immune exhaustion gene *TIGIT* (p=0.034) in NAT-treated patients compared to NAT-naïve patients. These results suggest that NAT-induced upregulation of TME complement expression may lead to reduced immune exhaustion and immunosuppression in PDAC, thereby contributing to the improved survival in these patients (Figure 6C).

**Figure. 6.**
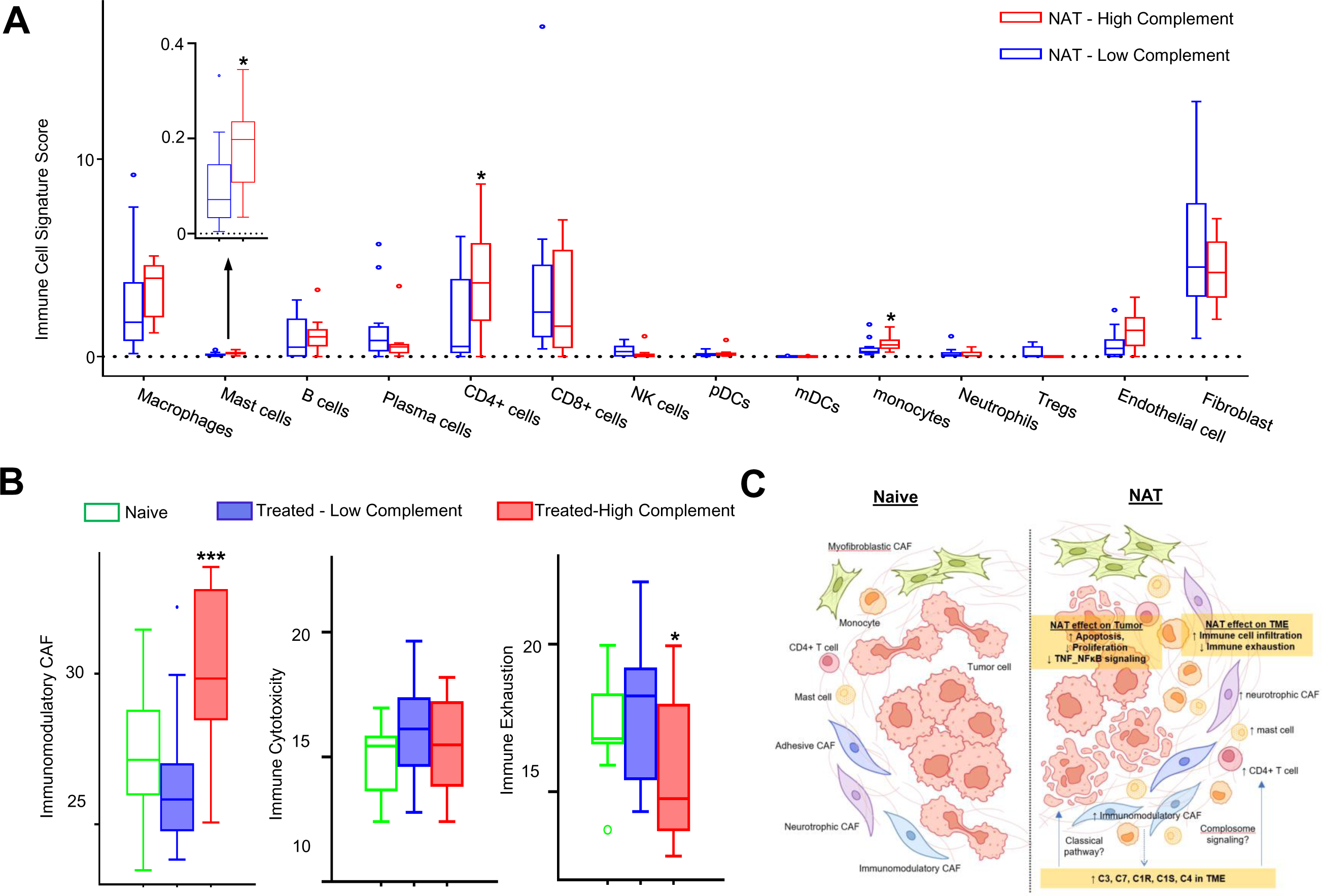
Association of NAT-induced upregulation of TME complement with immunomodulation and reduced immune exhaustion. **A.** Box-Whisker plot showing that the high TME complement subgroup has significantly higher levels of mast cells, CD4+ T cells and monocytes compared to the low TME complement subgroup (* p<0.05). **B.** The high TME complement subgroup has significantly increased immunomodulatory CAFs compared to the low TME complement group. In addition, the high TME complement subgroup has significantly decreased immune exhaustion signature score and no significant change in immune cytotoxicity signature score (* p<0.05; *** p<0.001). **C.** Schematic representation summarizing how NAT differentially remodels carcinoma cells and the TME of PDAC. While NAT mainly increases apoptosis and decreases the proliferation and G2/M phase cycling in carcinoma cells, it has broad immunomodulatory effects on the TME through upregulating local complement expression and signaling.

## Discussion

Our results revealed that NAT differentially remodels the carcinoma cells and the TME of PDAC through distinct mechanisms. By comprehensively comparing the spatial transcriptomics profiles of over 1800 genes related to cancer biology, TME and immune responses between a sizable cohort of patients treated with NAT versus those who were NAT-naïve, we demonstrated that NAT induced 44 DEGs in carcinoma cells and 52 DEGs in the TME. Gene set enrichment analysis revealed 9 significantly up-or downregulated gene sets and functional terms in NAT-treated carcinoma cells, compared to 61 in NAT-treated TME. These results indicate that NAT may exert a more extensive and profound remodeling effect on the TME than on the carcinoma cells per se. In the NAT-treated carcinoma cells, most DEGs are associated with apoptosis regulation, cell cycle control, and DNA repair. Among them, the most significantly upregulated DEG was *DDB2* (p=0.00077), which encodes DNA damage-binding protein 2, a nuclear protein crucial for DNA repair and genomic stability. Additionally, DDB2 has been shown to suppress epithelial-to-mesenchymal transition (EMT) and enhance sensitivity to chemotherapy in PDAC(42)^2^. Conversely, in the TME of NAT-treated PDAC, most DEGs are associated with complement signaling and regulation of innate and adaptive immune function. Among them, the most significant downregulated DEG was *HDAC2*, which encodes histone deacetylase 2, a transcription factor capable of inducing inflammatory gene expression in CAFs and supporting carcinoma growth in PDAC. Recent studies have demonstrated the role of HDAC2 in facilitating pancreatic cancer metastasis(43).

Notably, we observed a coordinated upregulation of five key complement genes (C3, C4B, C7, C1R, C1S) in the TME of NAT-treated PDAC as compared to those not treated by NAT. Due to the collective upregulation of these five complement genes, the complement pathway emerged as the most significantly enriched pathway (FDR=1.04×10^-5^) among the NAT upregulated gene sets. It is worth emphasizing that this NAT-induced upregulation of complement expression was spatially confined to TME. Traditionally, complement proteins were primarily considered to be produced in the liver and circulating in the bloodstream to aid the immune function by opsonization(44). However, our results suggest that the complements may also be produced locally within TME and play an important role in regulating immune response. Analysis of snRNA-seq data validated CAFs as the primary mediators of this NAT-induced upregulation of complement in TME, while complement gene expression in carcinoma cells and immune cells were not notably reduced. As fibroblasts are the most abundant cell type in TME, our results suggest that CAFs are likely the principal factor contributing to the local production of complement proteins within PDAC TME.

Our findings align with several recent single-cell or single-nucleus RNA-sequencing (sc/snRNA-Seq) studies. In 2021, Chen et al. identified a distinctive subtype of CAF, which they named “complement-secreting CAFs” (cs-CAFs), characterized by their enrichment with complement system genes. They found that cs-CAFs were predominantly located near malignant glands, and mainly existed in early-stage PDAC and diminished in late-stage PDAC(45). More recently, Croft et al. demonstrated that some CAFs located farther from malignant glands can also produce complement proteins, and their increase was associated with improved patient survival (46). In 2022, Hwang et al. (24) expanded the CAF classification system to include four subtypes: myofibroblastic, adhesive, immunomodulatory, and neurotrophic. They noted an overlap between the latter three subtypes and the previously defined inflammatory type CAFs, with immunomodulatory CAFs showing enrichment in complement genes (24).

Similarly, Werba et al. also observed complement gene enrichment in inflammatory CAFs (33). However, none of these sc/snRNA-Seq studies specifically investigated the impact of NAT on complement gene expression within TME. Our research, for the first time, demonstrated a NAT-induced upregulation of TME complement in CAFs. Further studies, especially those using single-cell spatial transcriptomics, are warranted to characterize how NAT influences the spatial distribution and adaptability of different subtypes of CAFs. A recent study by Shiau et al. using single cell spatial transcriptomics revealed that NAT induced significant changes in interaction strength between malignant cells and inflammatory CAFs mediated by enriched IL-6ST signaling (30). Correspondingly, our study also showed a significant upregulation of IL-6ST expression in post-NAT TME, contributing to the regulation of immune function (Figure 3A and 3C). Taken together, these results suggest that NAT can substantially remodel the TME and influence the interplay between carcinoma cells and inflammatory CAFs.

As another interesting finding of our study, NAT-induced upregulation of complement expression in TME was found to be associated with modulation of the TIME and decreased expression of immune exhaustion markers, which was demonstrated in both our spatial transcriptomics analysis (Figure 6B) and the snRNA-seq data. Importantly, the NAT-treated patients with higher TME complement demonstrated improved overall survival (Figure 5B). The detailed cellular and molecular mechanisms underlying these observations are yet to be further clarified. It is known that some cleaved complement proteins (such as C3a, C5a, etc.) are potent chemoattractant and can recruit immune cells into TIME(47–49). Recent evidence also indicated that a tonic level of complements in the microenvironment is required for maintaining immune cell homeostasis and function via the non-canonical intracellular complement (i.e., complosome) signaling (50,51). Our data suggests that TME complement may serve as a regulatory hub in the local tumor microenvironment for both innate and adaptive immune functions via both canonical and non-canonical signaling mechanisms. Further studies are warranted to elucidate the specific details of this novel mechanism.

Our study is the first spatial transcriptomics study specifically aimed at comparing the differential remodeling effect of NAT on carcinoma cells and the TME using FFPE tissue of a relatively large cohort of resected primary PDAC. The main limitation of our approach is that the spatial transcriptomics method used here does not provide single-cell resolution. On the other hand, since there is no cell isolation and processing needed as in single-cell or single-nucleus RNA-seq, the method used in this study reduces technical variability and batch effect, and is not impacted by the artifactual transcriptomics changes due to the procedure-induced cell stress. This is especially important for PDAC in which all cellular components are buried in excessive dense extracellular matrix which makes cell isolation particularly challenging. In addition, by pooling mRNAs from each AOI, this method can achieve higher mRNA concentration per sample, thus facilitating deeper sequencing and reduced dropouts, thereby enhancing the signal-to-noise ratio. Consequently, it enables more robust detection of weakly expressed genes. Recently, the scRNA-seq study by Werba et al. observed generally decreased cell-cell interaction and high volume of putative interactions between CAFs and tumor-associated macrophages (TAMs) in the TME of NAT-treated PDAC (33). No difference was noted in CAF subpopulation distributions between NAT-treated vs. naive groups(33). Hwang et al. discovered that NAT could increase the adhesive CAFs and decrease the myofibroblast CAFs. They also found that the number of immunomodulatory CAFs was higher in patients treated with regular NAT than those with NAT plus losartan (24). In the most recent study using single-cell spatial transcriptomics, Shiau et al showed a marked increased ligand-receptor interaction between CAFs and malignant cells in NAT-treated PDAC (30). Therefore, our data have not only confirmed previous sc/snRNA-seq studies but also provided additional valuable insights thanks to the preserved spatial context.

In conclusion, by using spatially resolved transcriptomics, we demonstrated that NAT remodels the carcinoma cells and the TME of PDACs through different mechanisms. NAT can significantly upregulate complement gene expression within TME, which is associated with reduced immune exhaustion and improved survival. These results suggest that NAT may potentially attenuate immune cell exhaustion and enhance immune surveillance in some PDAC patients by upregulating local complement production and signaling. Those patients who did not respond to NAT with a significant upregulation of TME complement may have more immune exhaustion and thus benefit from combinational immune checkpoint blockade therapy. Therefore, assessment of the local complement dynamics in the TME may provide more accurate prognostic guidance for NAT-treated patients, which may help to stratify them for more tailored downstream interventions. Further analyses, especially single-cell spatial transcriptomics with intact spatial context, are warranted to further elucidate the detailed cellular and molecular mechanisms of complement-mediated immunomodulation and its potential therapeutic implications.

## Author contributions

X.Z., Y.Z.L., and Y.L. developed the study concept and the study design. X.Z., Y.Z.L., Y.L., and R.L. wrote the manuscript, which was then edited by all co-authors. X.Z. acquired slides and tissue blocks, collected patient information, and helped with the GeoMx experiment. X.Z. and D.L. analyzed the H&E and multiplex IHC. R.L., Y.Z.L, and X.Z. analyzed spatial transcriptomics data and conducted statistical analysis of clinicopathological information. V.G.P. provided feedback on the study design and the manuscript. C.L.Z., S.A.S., W.L., I.H., M.G., C.H., J.M., J.W., M.S, J.A., and M.K. provided feedback and edited the manuscript. All co-authors approved the final version of the manuscript before submission.

## Acknowledgement

The authors thank the University of Wisconsin Translational Research Initiatives in Pathology laboratory (TRIP), supported by the UW Department of Pathology and Laboratory Medicine, UWCCC (P30 CA014520) and the Office of The Director-NIH (S10 OD023526) for use of its facilities and services. The authors also thank Nancy Liu for the artistic illustration in Figure 6C summarizing the main findings of this study.

## Supplementary Figures

**Supplementary Figure 1.**
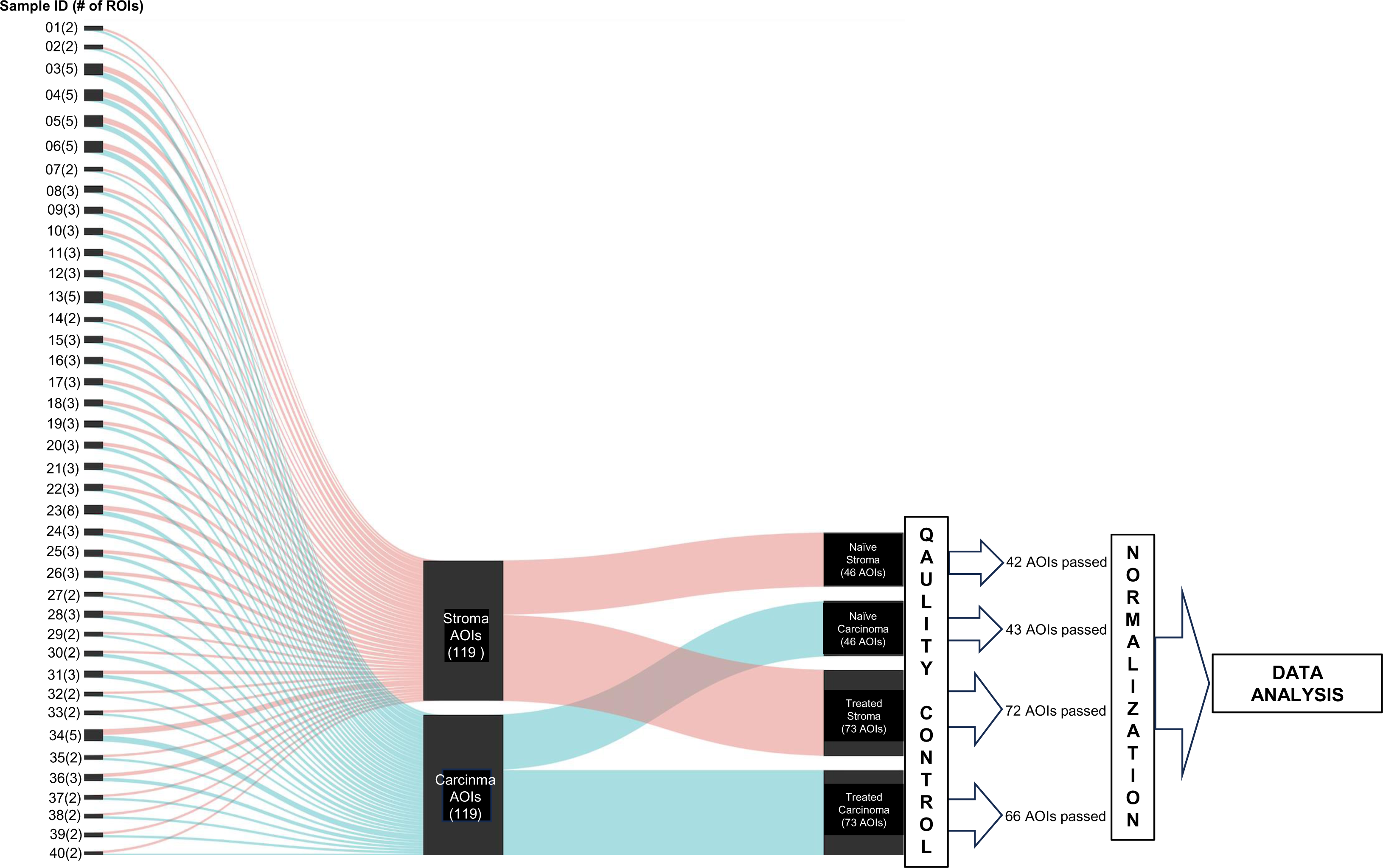
The overall experimental Design.

**Supplementary Figure 2.**
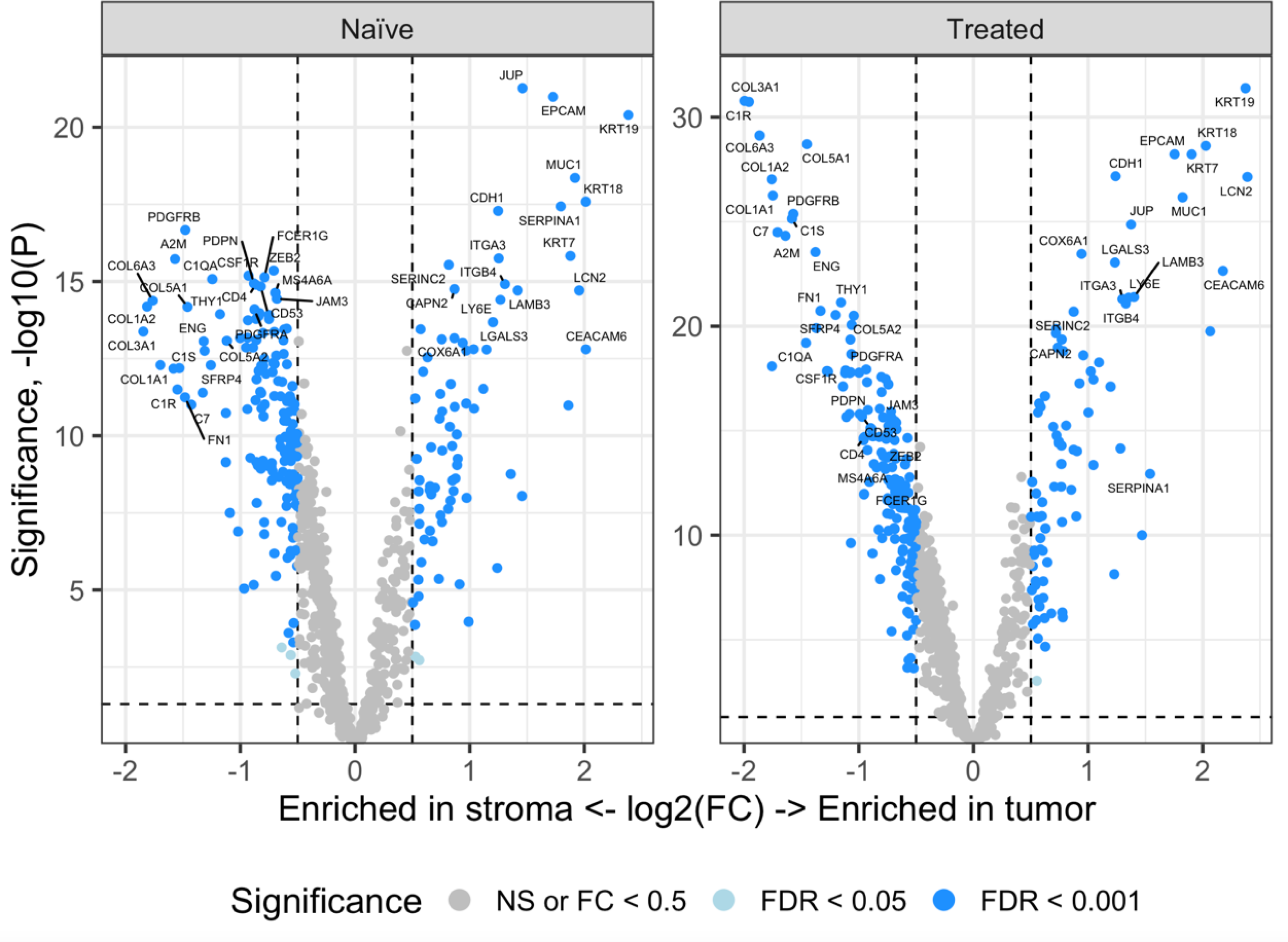
The volcano plots of DEGs in carcinoma AOIs and TME AOIs in the naïve group and the NAT group.

**Supplementary Figure 3.**
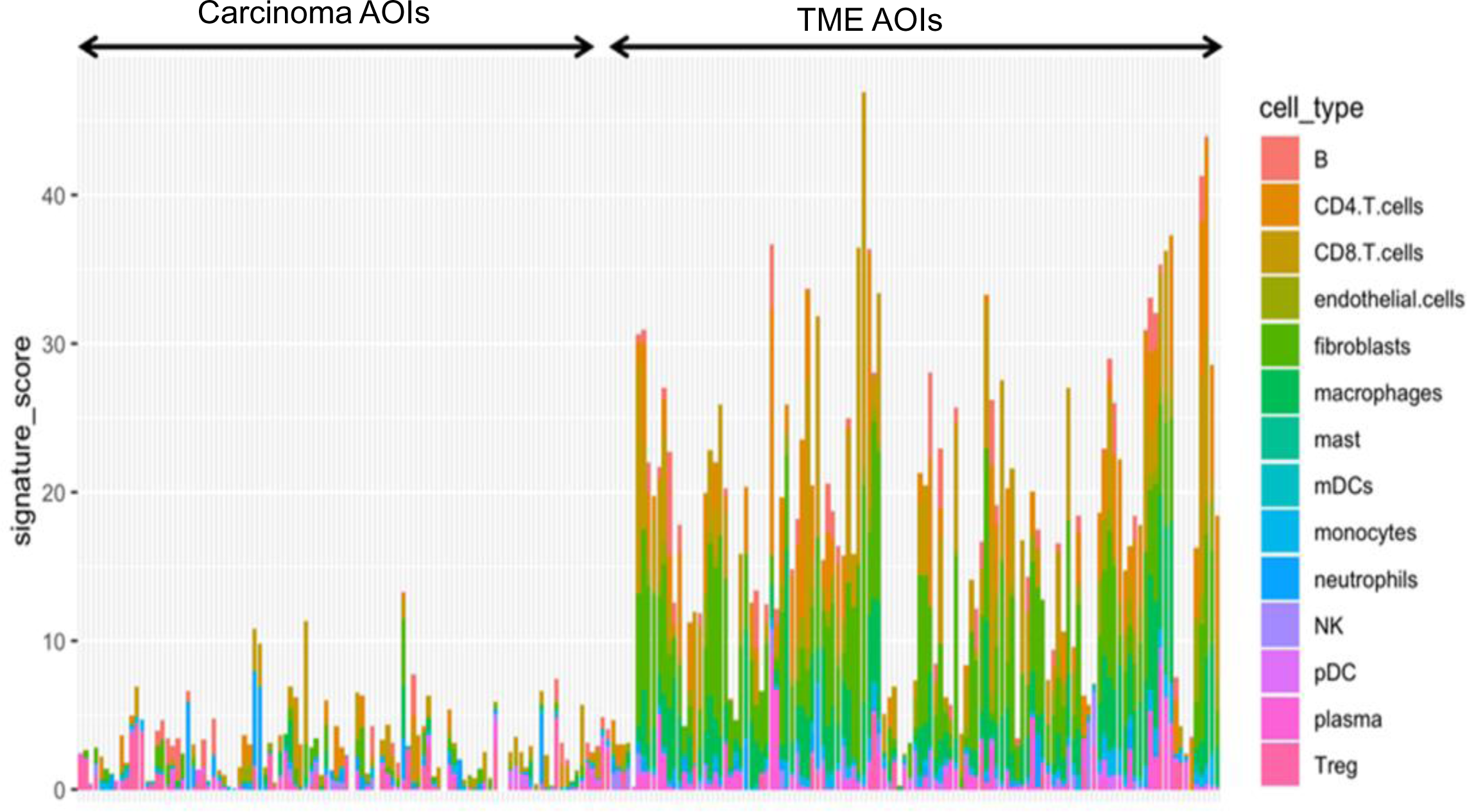
Immune cell distribution in carcinoma and TME AOIs across the study cohort. The TME AOIs exhibit significantly greater abundance and diversity of immune cells compared to the carcinoma AOIs.

**Supplementary Figure 4.**
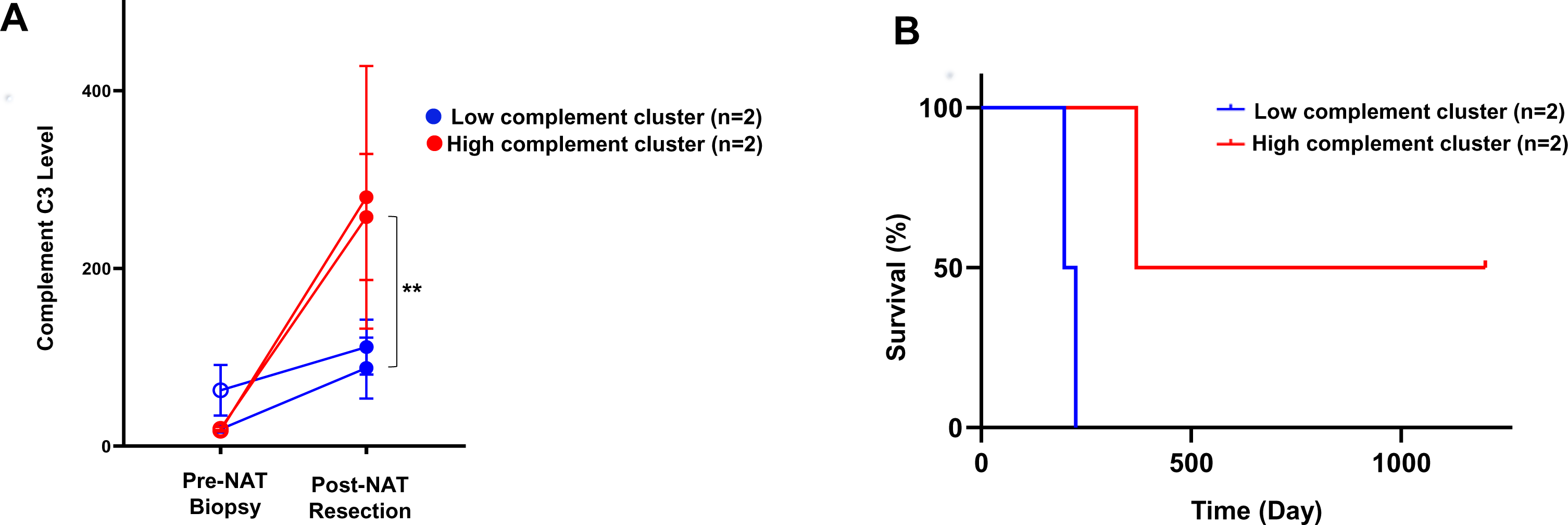
Paired analysis of pre-NAT biopsy and post-NAT resection from the same patient. **A.** Among the four patients for whom both the pre-NAT biopsy and post-NAT resection were available, two of them exhibited NA-induced upregulation of C3 complement level in TME, while the other two patients showed no increase. **B.** The two patients with NAT-induced upregulation of TME C3 complement level showed better overall survival.

**Supplementary Table. 1.** Clinicopathologic characteristics of the NAT-naïve patients and NAT-treated patients in our study cohort.

| Characteristics | All subjects | NAT group | Naïve group |
| --- | --- | --- | --- |
| Total (n) | 36 | 23 | 13 |
| Median age, years (range) | 65.5 (49-86) | 66 (59-78) | 63 (49-86) |
| Age group (years), n (%) |  |  |  |
| <65 | 15 (41.7) | 8 (34.8) | 7 (53.8) |
| >=65 | 21 (58.3) | 15 (65.2) | 6 (46.2) |
| Gender, n (%) |  |  |  |
| Male | 17 (47.2) | 9 (39.1) | 8 (61.5) |
| Female | 19 (52.8) | 14 (60.9) | 5 (38.5) |
| Tumor size (range) |  |  |  |
| 0.5 – 2.0 | 9 (25.0) | 7 (30.4) | 2 (15.4) |
| 2.1 – 4.0 | 20 (55.6) | 12 (52.2) | 8 (61.5) |
| >4.0 | 7 (19.4) | 4 (17.4) | 3 (23.1) |
| Cancer grading, n (%) |  |  |  |
| G1 | 6 (16.7) | 3 (13.0) | 3 (23.1) |
| G2 | 20 (55.6) | 12 (52.2) | 8 (61.5) |
| G3 | 10 (27.7) | 8 (34.8) | 2 (15.4) |
| Staging, n (%) |  |  |  |
| I and II | 23 (63.9) | 17 (73.9) | 6 (46.2) |
| III and IV | 13 (36.1) | 6 (26.1) | 7 (53.8) |

**Supplementary Table. 2.** Clinicopathologic characteristics of 4 patients for the comparison between pre-NAT biopsy and post-NAT resection.

| Characteristics |  | All subjects | High TME C3 group | Low TME C3 group |
| --- | --- | --- | --- | --- |
| Total, n |  | 4 | 2 | 2 |
| Median age, years (range) |  | 69.5 (67-73) | 68 (67-69) | 71.5 (70-73) |
| Age group (years), n (%) |  |  |  |  |
|  | <70 | 2 (50.0) | 2 (100.0) | 0 (0) |
|  | >=70 | 2 (50.0) | 0 (0) | 2 (100.0) |
| Gender, n (%) |  |  |  |  |
|  | Male | 1 (25.0) | 0 (0) | 1 (50.0) |
|  | Female | 3 (75.0) | 2 (100.0) | 1 (50.0) |
| Tumor size, cm (%) |  |  |  |  |
|  | 0.5 – 2.0 | 1 (25.0) | 0 (0) | 1 (50.0) |
|  | 2.1 – 4.0 | 2 (50.0) | 2 (100.0) | 0 (0) |
|  | >4.0 | 1 (25.0) | 0 (0) | 1 (50.0) |
| Cancer grading, n (%) |  |  |  |  |
|  | G1 | 0 (0) | 0 (0) | 0 (0) |
|  | G2 | 3 (75.0) | 2 (100.0) | 1 (50.0) |
|  | G3 | 1 (25.0) | 0 (0) | 1 (50.0) |
| Staging, n (%) |  |  |  |  |
|  | I | 1 (25) | 1 (25) | 1 (25) |
|  | II | 3 (75) | 3 (75) | 3 (75) |

## References

1. Siegel RL, Miller KD, Fuchs HE, Jemal A. Cancer statistics, 2022. CA Cancer J Clin 2022;72(1):7-33 doi 10.3322/caac.21708.

2. Rahib L, Wehner MR, Matrisian LM, Nead KT. Estimated Projection of US Cancer Incidence and Death to 2040. JAMA Netw Open 2021;4(4):e214708 doi 10.1001/jamanetworkopen.2021.4708.

3. Vincent A, Herman J, Schulick R, Hruban RH, Goggins M. Pancreatic cancer. Lancet 2011;378(9791):607-20 doi 10.1016/S0140-6736(10)62307-0.

4. Ferrone CR, Brennan MF, Gonen M, Coit DG, Fong Y, Chung S, et al. Pancreatic adenocarcinoma: the actual 5-year survivors. J Gastrointest Surg 2008;12(4):701–6 doi 10.1007/s11605-007-0384-8.

5. Wood LD, Canto MI, Jaffee EM, Simeone DM. Pancreatic Cancer: Pathogenesis, Screening, Diagnosis, and Treatment. Gastroenterology 2022;163(2):386–402 e1 doi 10.1053/j.gastro.2022.03.056.

6. Manrai M, Tilak T, Dawra S, Srivastava S, Singh A. Current and emerging therapeutic strategies in pancreatic cancer: Challenges and opportunities. World J Gastroenterol 2021;27(39):6572–89 doi 10.3748/wjg.v27.i39.6572.

7. Collisson EA, Bailey P, Chang DK, Biankin AV. Molecular subtypes of pancreatic cancer. Nat Rev Gastroenterol Hepatol 2019;16(4):207–20 doi 10.1038/s41575-019-0109-y.

8. Herting CJ, Karpovsky I, Lesinski GB. The tumor microenvironment in pancreatic ductal adenocarcinoma: current perspectives and future directions. Cancer Metastasis Rev 2021;40(3):675–89 doi 10.1007/s10555-021-09988-w.

9. Shibuya KC, Goel VK, Xiong W, Sham JG, Pollack SM, Leahy AM, et al. Pancreatic ductal adenocarcinoma contains an effector and regulatory immune cell infiltrate that is altered by multimodal neoadjuvant treatment. PLoS One 2014;9(5):e96565 doi 10.1371/journal.pone.0096565.

10. Hosein AN, Brekken RA, Maitra A. Pancreatic cancer stroma: an update on therapeutic targeting strategies. Nat Rev Gastroenterol Hepatol 2020;17(8):487–505 doi 10.1038/s41575-020-0300-1.

11. Su H, Yang F, Fu R, Trinh B, Sun N, Liu J, et al. Collagenolysis-dependent DDR1 signalling dictates pancreatic cancer outcome. Nature 2022;610(7931):366-72 doi 10.1038/s41586-022-05169-z.

12. Murphy JE, Wo JY, Ryan DP, Jiang W, Yeap BY, Drapek LC, et al. Total Neoadjuvant Therapy With FOLFIRINOX Followed by Individualized Chemoradiotherapy for Borderline Resectable Pancreatic Adenocarcinoma: A Phase 2 Clinical Trial. JAMA Oncol 2018;4(7):963–9 doi 10.1001/jamaoncol.2018.0329.

13. Versteijne E, van Dam JL, Suker M, Janssen QP, Groothuis K, Akkermans-Vogelaar JM, et al. Neoadjuvant Chemoradiotherapy Versus Upfront Surgery for Resectable and Borderline Resectable Pancreatic Cancer: Long-Term Results of the Dutch Randomized PREOPANC Trial. J Clin Oncol 2022;40(11):1220–30 doi 10.1200/JCO.21.02233.

14. Attaallah W. Neoadjuvant Chemoradiotherapy for Resectable and Borderline Resectable Pancreatic Cancer. J Clin Oncol 2022;40(28):3346 doi 10.1200/JCO.22.00432.

15. Krishnamoorthy M, Lenehan JG, Maleki Vareki S. Neoadjuvant Immunotherapy for High-Risk, Resectable Malignancies: Scientific Rationale and Clinical Challenges. J Natl Cancer Inst 2021;113(7):823–32 doi 10.1093/jnci/djaa216.

16. Patel RB, Hernandez R, Carlson P, Grudzinski J, Bates AM, Jagodinsky JC, et al. Low-dose targeted radionuclide therapy renders immunologically cold tumors responsive to immune checkpoint blockade. Sci Transl Med 2021;13(602) doi 10.1126/scitranslmed.abb3631.

17. Kuipers J, Jahn K, Beerenwinkel N. Advances in understanding tumour evolution through single-cell sequencing. Biochim Biophys Acta Rev Cancer 2017;1867(2):127–38 doi 10.1016/j.bbcan.2017.02.001.

18. Lahnemann D, Koster J, Szczurek E, McCarthy DJ, Hicks SC, Robinson MD, et al. Eleven grand challenges in single-cell data science. Genome Biol 2020;21(1):31 doi 10.1186/s13059-020-1926-6.

19. Ali LR, Lenehan PJ, Cardot-Ruffino V, Dias Costa A, Katz MHG, Bauer TW, et al. PD-1 blockade induces reactivation of non-productive T cell responses characterized by NF-kB signaling in patients with pancreatic cancer. Clin Cancer Res 2023 doi 10.1158/1078-0432.CCR-23-1444.

20. Barthel S, Falcomata C, Rad R, Theis FJ, Saur D. Single-cell profiling to explore pancreatic cancer heterogeneity, plasticity and response to therapy. Nat Cancer 2023;4(4):454–67 doi 10.1038/s43018-023-00526-x.

21. Chen K, Ma Y, Liu X, Zhong X, Long D, Tian X, et al. Single-cell RNA-seq reveals characteristics in tumor microenvironment of PDAC with MSI-H following neoadjuvant chemotherapy with anti-PD-1 therapy. Cancer Lett 2023;576:216421 doi 10.1016/j.canlet.2023.216421.

22. Cui Zhou D, Jayasinghe RG, Chen S, Herndon JM, Iglesia MD, Navale P, et al. Spatially restricted drivers and transitional cell populations cooperate with the microenvironment in untreated and chemo-resistant pancreatic cancer. Nat Genet 2022;54(9):1390–405 doi 10.1038/s41588-022-01157-1.

23. Han J, DePinho RA, Maitra A. Single-cell RNA sequencing in pancreatic cancer. Nat Rev Gastroenterol Hepatol 2021;18(7):451–2 doi 10.1038/s41575-021-00471-z.

24. Hwang WL, Jagadeesh KA, Guo JA, Hoffman HI, Yadollahpour P, Reeves JW, et al. Single-nucleus and spatial transcriptome profiling of pancreatic cancer identifies multicellular dynamics associated with neoadjuvant treatment. Nat Genet 2022;54(8):1178–91 doi 10.1038/s41588-022-01134-8.

25. Kinker GS, Vitiello GAF, Diniz AB, Cabral-Piccin MP, Pereira PHB, Carvalho MLR, et al. Mature tertiary lymphoid structures are key niches of tumour-specific immune responses in pancreatic ductal adenocarcinomas. Gut 2023;72(10):1927–41 doi 10.1136/gutjnl-2022-328697.

26. Li K, Tandurella JA, Gai J, Zhu Q, Lim SJ, Thomas DL, 2nd, et al. Multi-omic analyses of changes in the tumor microenvironment of pancreatic adenocarcinoma following neoadjuvant treatment with anti-PD-1 therapy. Cancer Cell 2022;40(11):1374-91 e7 doi 10.1016/j.ccell.2022.10.001.

27. Lin W, Noel P, Borazanci EH, Lee J, Amini A, Han IW, et al. Single-cell transcriptome analysis of tumor and stromal compartments of pancreatic ductal adenocarcinoma primary tumors and metastatic lesions. Genome Med 2020;12(1):80 doi 10.1186/s13073-020-00776-9.

28. Peng J, Sun BF, Chen CY, Zhou JY, Chen YS, Chen H, et al. Single-cell RNA-seq highlights intra-tumoral heterogeneity and malignant progression in pancreatic ductal adenocarcinoma. Cell Res 2019;29(9):725–38 doi 10.1038/s41422-019-0195-y.

29. Raghavan S, Winter PS, Navia AW, Williams HL, DenAdel A, Lowder KE, et al. Microenvironment drives cell state, plasticity, and drug response in pancreatic cancer. Cell 2021;184(25):6119–37 e26 doi 10.1016/j.cell.2021.11.017.

30. Shiau C, Cao J, Gregory MT, Gong D, Yin X, Cho JW, et al. Therapy-associated remodeling of pancreatic cancer revealed by single-cell spatial transcriptomics and optimal transport analysis. bioRxiv 2023 doi 10.1101/2023.06.28.546848.

31. Shiau C, Su J, Guo JA, Hong TS, Wo JY, Jagadeesh KA, et al. Treatment-associated remodeling of the pancreatic cancer endothelium at single-cell resolution. Front Oncol 2022;12:929950 doi 10.3389/fonc.2022.929950.

32. Ting DT, Wittner BS, Ligorio M, Vincent Jordan N, Shah AM, Miyamoto DT, et al. Single-cell RNA sequencing identifies extracellular matrix gene expression by pancreatic circulating tumor cells. Cell Rep 2014;8(6):1905–18 doi 10.1016/j.celrep.2014.08.029.

33. Werba G, Weissinger D, Kawaler EA, Zhao E, Kalfakakou D, Dhara S, et al. Single-cell RNA sequencing reveals the effects of chemotherapy on human pancreatic adenocarcinoma and its tumor microenvironment. Nat Commun 2023;14(1):797 doi 10.1038/s41467-023-36296-4.

34. Hou Z, Lin J, Ma Y, Fang H, Wu Y, Chen Z, et al. Single-cell RNA sequencing revealed subclonal heterogeneity and gene signatures of gemcitabine sensitivity in pancreatic cancer. Front Pharmacol 2023;14:1193791 doi 10.3389/fphar.2023.1193791.

35. Yousuf S, Qiu M, Voith von Voithenberg L, Hulkkonen J, Macinkovic I, Schulz AR, et al. Spatially Resolved Multi-Omics Single-Cell Analyses Inform Mechanisms of Immune Dysfunction in Pancreatic Cancer. Gastroenterology 2023;165(4):891–908 e14 doi 10.1053/j.gastro.2023.05.036.

36. Longo SK, Guo MG, Ji AL, Khavari PA. Integrating single-cell and spatial transcriptomics to elucidate intercellular tissue dynamics. Nat Rev Genet 2021;22(10):627–44 doi 10.1038/s41576-021-00370-8.

37. Rao A, Barkley D, Franca GS, Yanai I. Exploring tissue architecture using spatial transcriptomics. Nature 2021;596(7871):211-20 doi 10.1038/s41586-021-03634-9.

38. Huang da W, Sherman BT, Lempicki RA. Systematic and integrative analysis of large gene lists using DAVID bioinformatics resources. Nat Protoc 2009;4(1):44–57 doi 10.1038/nprot.2008.211.

39. Sherman BT, Hao M, Qiu J, Jiao X, Baseler MW, Lane HC, et al. DAVID: a web server for functional enrichment analysis and functional annotation of gene lists (2021 update). Nucleic Acids Res 2022;50(W1):W216–W21 doi 10.1093/nar/gkac194.

40. Hao Y, Stuart T, Kowalski MH, Choudhary S, Hoffman P, Hartman A, et al. Dictionary learning for integrative, multimodal and scalable single-cell analysis. Nat Biotechnol 2023 doi 10.1038/s41587-023-01767-y.

41. Satija R, Farrell JA, Gennert D, Schier AF, Regev A. Spatial reconstruction of single-cell gene expression data. Nat Biotechnol 2015;33(5):495–502 doi 10.1038/nbt.3192.

42. Dardare J, Witz A, Betz M, Francois A, Meras M, Lamy L, et al. DDB2 represses epithelial-to-mesenchymal transition and sensitizes pancreatic ductal adenocarcinoma cells to chemotherapy. Front Oncol 2022;12:1052163 doi 10.3389/fonc.2022.1052163.

43. Krauss L, Urban BC, Hastreiter S, Schneider C, Wenzel P, Hassan Z, et al. HDAC2 Facilitates Pancreatic Cancer Metastasis. Cancer Res 2022;82(4):695–707 doi 10.1158/0008-5472.CAN-20-3209.

44. Merle NS, Noe R, Halbwachs-Mecarelli L, Fremeaux-Bacchi V, Roumenina LT. Complement System Part II: Role in Immunity. Front Immunol 2015;6:257 doi 10.3389/fimmu.2015.00257.

45. Chen K, Wang Q, Li M, Guo H, Liu W, Wang F, et al. Single-cell RNA-seq reveals dynamic change in tumor microenvironment during pancreatic ductal adenocarcinoma malignant progression. EBioMedicine 2021;66:103315 doi 10.1016/j.ebiom.2021.103315.

46. Croft W, Pearce H, Margielewska-Davies S, Lim L, Nicol SM, Zayou F, et al. Spatial determination and prognostic impact of the fibroblast transcriptome in pancreatic ductal adenocarcinoma. Elife 2023;12 doi 10.7554/eLife.86125.

47. Ajona D, Ortiz-Espinosa S, Pio R, Lecanda F. Complement in Metastasis: A Comp in the Camp. Front Immunol 2019;10:669 doi 10.3389/fimmu.2019.00669.

48. Kolev M, Das M, Gerber M, Baver S, Deschatelets P, Markiewski MM. Inside-Out of Complement in Cancer. Front Immunol 2022;13:931273 doi 10.3389/fimmu.2022.931273.

49. Pio R, Ajona D, Ortiz-Espinosa S, Mantovani A, Lambris JD. Complementing the Cancer-Immunity Cycle. Front Immunol 2019;10:774 doi 10.3389/fimmu.2019.00774.

50. West EE, Kemper C. Complosome - the intracellular complement system. Nat Rev Nephrol 2023;19(7):426–39 doi 10.1038/s41581-023-00704-1.

51. Shao F, Jin K, Li B, Liu Z, Zeng H, Wang Y, et al. Integrating angiogenesis signature and tumor mutation burden for improved patient stratification in immune checkpoint blockade therapy for muscle-invasive bladder cancer. Urol Oncol 2023;41(10):433 e9-e18 doi 10.1016/j.urolonc.2023.07.006.

